# Alternative reproductive morphs and sexes differ in DNA methylation along distinct genomic axes in *Poecilia parae*

**DOI:** 10.64898/2026.11.25.769244

**Authors:** Maximiliano Zuluaga-Forero, Ehren Bentz, Ben Sandkam

## Abstract

Biological diversity occurs not only between species and sexes, but also within the sexes. Differences between males and females (intersexual differences) are well known to correspond to distinct DNA methylation differences, suggesting that differences between alternative phenotypes of the same sex (intrasexual differences) may also correspond to such methylation differences. Here, we used the fish *Poecilia parae*, in which males occur as discrete Y-linked morphs, to ask whether heritable diversity within the sexes corresponds to alternative DNA methylomes. Using whole-genome enzymatic methyl-seq of adult tail muscle from individuals of three male morphs and females, we quantified intersexual and intrasexual methylation divergence. Intrasexual differences in methylation were distributed broadly across the shared autosomes, without a skew toward extreme methylation, whereas there were strong intersexual differences across the X chromosome. Morph-specific regions of differential methylation recurred across contrasts modestly but consistently more often than in randomized groupings of the same individuals, on both the autosomes and the X chromosome, indicating that alternative male morphs carry distinguishable methylation profiles. These results show that a heritable within-sex polymorphism can be reflected in alternative epigenomic states across a shared genomic background.

**SIGNIFICANCE STATEMENT:** Within species, DNA methylation profiles differ between the sexes and are thought to play a major role in mediating sexually dimorphic traits. In parallel, intraspecific diversity expressed within a sex may also correspond to alternative methylomes. By comparing methylation state between discrete alternative male morphs of the fish *Poecilia parae* we found that alternative reproductive phenotypes within a sex also correspond to alternative genome-wide methylation profiles. These intrasexual differences suggest that methylation profiles can differ among heritable phenotypes within a sex and may contribute to phenotypic diversity within sexes.

## INTRODUCTION

Biological diversity occurs at multiple levels of organization; there are often differences in morphology, physiology, and behavior across species, between sexes, and even among individuals of the same species and sex (Rubenstein 1980; West-Eberhard 1986; Oliveira et al. 2008). Although differences between species often involve numerous genetic changes accumulated across the genome, diversity within species can arise from comparatively limited genetic differences, such as a sex-determining chromosome underlying sex differences (Bull 1983; Charlesworth 1996), or even a restricted genomic region specifying alternative phenotypes (Fisher 1930; Schwander et al. 2014). Despite their relatively small genomic footprint, these regions mediating heritable diversity are often associated with broad differences in gene expression across the genome (Clarke and Sheppard 1960; Charlesworth 1996), suggesting that intraspecific diversity may correspond with genome-wide differences in epigenetic states, such as CpG methylation.

CpG methylation is one of the most widespread and well-characterized epigenetic modifications in vertebrates. In the same species and tissue, genome-wide methylation profiles are broadly reproducible among individuals, often functioning as molecular signatures of cellular identity (Jaenisch and Bird 2003; Rakyan et al. 2004; Eckhardt et al. 2006; Lister et al. 2009; Smith et al. 2015; Bogdanović et al. 2016; Al Adhami et al. 2022). However, even within the same tissue, well-documented sources of methylome differentiation still exist, such as biological sex (Liu et al. 2010; Duncan et al. 2018; Metzger and Schulte 2018; Metzger and Mank 2026). These sex-associated differences are accompanied by striking differences in gene expression (Rinn and Snyder 2005; Metzger et al. 2021; Darolti et al. 2023), suggesting that sex-associated methylation differences may reflect biologically meaningful molecular differentiation and positioning biological sex as one of the clearest examples of methylome diversity within species.

Sex-associated differences in methylation have been documented across vertebrates, including mammals and fish, often at modest magnitudes (Liu et al. 2010; Sun et al. 2016; Metzger and Schulte 2018). For example, genome-wide intersexual differentiation has been estimated at approximately 0.7% in tilapia (Sun et al. 2016) and ∼1% in guppies (Metzger and Mank 2026). Importantly, even such magnitudes of methylation divergence can be sufficiently distinct to classify an individual’s sex (Wang et al. 2021). Although sex-associated methylation differences are often more pronounced on sex chromosomes than on autosomes, they also occur across the autosomal genome. In humans, autosomal methylation differs between the sexes by approximately 0.9% (McCarthy et al. 2014). Similarly, in somatic tissue of three-spined stickleback, approximately 0.07% of autosomal CpGs are differentially methylated between the sexes, compared with 5% on the sex chromosome (Metzger and Schulte 2018). Therefore, even modest autosomal methylation differences are also reproducible and can be associated with distinct sexual phenotypes (Liu et al. 2010; Thomson et al. 2017; White et al. 2017; Sandkam 2026).

Methylation divergence has also been observed in alternative phenotypes within the same sex (Metzger and Schulte 2018; Sun et al. 2021; Matlosz et al. 2022; Prichard et al. 2022). However, the magnitude and genomic distribution of such differentiation remain poorly understood. Understanding these patterns is especially relevant in species where individuals of the same sex occur as discrete, genetically inherited morphs (Sandkam 2026). These morphs often arise from relatively narrow regions of genetic differentiation but show widespread differences in gene expression across their shared genome, along with differences in morphology, physiology, behavior, and life history (West-Eberhard 1986; Oliveira et al. 2008). Because the autosome constitutes most of the shared genome, it provides a common genomic background across which extensive morph-associated epigenetic differences should occur, with CpG methylation well positioned to contribute to such alternatives over the shared sequence (De Mendoza et al. 2020; Klughammer et al. 2023). If methylation correlates with within-sex differences as it does with differences between sexes, then alternative morphs should exhibit methylation divergence across their shared autosomal genome at a magnitude comparable to intersexual autosomal divergence.

The polymorphic fish *Poecilia parae* offers an exceptional opportunity to test this hypothesis. Males of this species develop into one of three discrete major morphs that differ strikingly in body size, color patterns, and mating strategy. The Parae morph has the largest body and vertical black stripes, and they are more aggressive toward conspecifics. The Melanzona morph has a medium-sized body with a horizontal colored stripe (yellow, blue, or red), which they use to display to females. Meanwhile, the Immaculata morph resembles a juvenile female. They have the smallest body, but the largest testes (Liley 1966; Bourne et al. 2003; Hurtado-Gonzales and Uy 2009; Stec et al. 2026). Morph identity is inherited strictly from father to son through differences only on the Y chromosome, while all morphs share the same X chromosome and the autosomes (Lindholm et al. 2004; Sandkam et al. 2021). Differences in gene regulation have been shown to occur between the sexes across the entire X chromosome of *P. parae* (Metzger et al. 2021), as well as in its close relatives (*Poecilia picta* (Darolti et al. 2019) and *Poecilia bifurca* (Fong et al. 2025). Additionally, a recent study in the sister species *P. picta* found that these sex differences in X chromosome regulation are accompanied by strong sex differences in CpG methylation across the X chromosome, suggesting DNA methylation may play a role in broad regulatory differences between the sexes (Metzger et al. 2023).

Here, we ask whether the alternative male morphs of *P. parae* are reflected in alternative CpG methylation profiles across the genome. We generated enzymatic methylation sequencing data across the whole genome of each male morph and females. We quantified methylation divergence at both single-CpG and regional scales, with particular emphasis on the autosomes, where the sequence background is shared. By comparing inter to intrasexual methylation divergence across the autosomes and the X chromosome, we assessed the magnitude of biologically meaningful DNA methylation differences across phenotypes. To distinguish phenotype-associated differences from individual variation, we tested whether regions of differential methylation recur in all contrasts involving a given phenotype more often than in contrasts between randomized groups of the same individuals. We found widespread autosomal methylation differentiation among male morphs, with recurrent patterns of regional differentiation associated with morph identity. Together, our findings indicate that differences in genome-wide methylomes reflect the alternative reproductive morphs of *P. parae*.

## Results

### Differential methylation is present across the whole genome between morphs and sexes

To test whether the alternative phenotypes are associated with overall differences in CpG methylation profiles, we ran enzymatic methylation sequencing (EMSeq) on three individuals from each of the three major-morphs of *Poecilia parae* (Immaculata, Parae, and Melanzona) and females **(Supplementary Figure 1)**. We then compared genome-wide methylation levels among females and the male morphs using Permutational Multivariate Analysis of Variance (PERMANOVA). Whole-genome CpG methylation profiles differed significantly among phenotypes (pseudo-F_3,8_ = 1.079, R^2^ = 0.288, p = 0.0229). With four groups of three individuals, R^2^ is expected to be ∼0.27 even in the absence of any phenotype effect, so the effect of phenotype on genome-wide methylation is small but consistent. This association was not explained by differences in within-phenotype variability, as Permutational Multivariate Analysis of Dispersion (PERMDISP) did not differ among phenotype groups (F_3,8_ = 0.048, p = 0.9812). Consistent with the PERMANOVA result, individuals belonging to the same phenotype had significantly smaller pairwise methylation differences than individuals belonging to different phenotypes (mean within-phenotype distance = 0.0897; mean between-phenotype distance = 0.0908; Delta = 0.00116; permutation p = 0.0334). Together, these results indicate that whole-genome CpG methylation shows modest but significant structuring by phenotype, with individuals of the same phenotype having more similar methylation profiles than individuals from different phenotypes.

### Single CpGs: morph differences are spread across the genome, whereas sex differences are enriched on the X chromosome

To identify the number of CpG sites where methylation differs between the sexes and morphs, we performed independent pairwise contrasts for each morph against females, as well as contrasts among the male morphs. Pairwise contrasts between females and each of the three male morphs identified 31,272 – 43,995 *Differentially Methylated Cytosines* (DMCs) across the genome (0.21 – 0.30%). Surprisingly, this was very similar to the number of DMCs identified in pairwise contrasts between the male-morphs, with Parae-vs-Immaculata having 40,651 (0.27%) DMCs, and Parae-vs-Melanzona having 42,299 (0.29%) DMCs. However, the Melanzona and Immaculata morphs were much more similar to one another, having only 28,915 (0.19%) DMCs between them **(Supplementary Table 1).**

The DMCs between males and females were not evenly distributed across the genome; instead, the proportion of differentially methylated CpG sites was roughly 10 times higher on the X chromosome (1.07–1.13% CpG sites) compared to the autosomes (0.17–0.26% CpG sites). However, this pattern did not hold when comparing methylation between the male morphs; instead, the percentage of DMCs was similar on the X chromosome (0.15–0.22% CpG sites) and the autosomes (0.20–0.29% CpG sites) **(Supplementary Table 1)**. These distributions are consistent with the previously proposed role of DNA methylation as a mechanism of X-chromosome dosage compensation, which is shared across all morphs (Darolti et al. 2019; Metzger et al. 2021; Metzger et al. 2023; Fong et al. 2025).

To visualize the genome-wide patterns of DMC distribution, we plotted DMC density in 100-kb windows across the genome. As expected, each of the morphs compared to females had elevated rates of DMCs across the entire X chromosome **(Figure 1)**. Meanwhile, DMCs between the male morphs were distributed across the whole genome, with occasional spikes in single pairwise contrasts (e.g. chromosomes 8, 11 and 21). The comparable numbers of DMCs in sex and male-morph contrasts and their widespread autosomal distribution suggest that methylation differentiation among male morphs may be comparable to differentiation between sexes across the autosomal genome, whereas sex-associated differences are particularly pronounced on the X chromosome.

**Figure 1.**
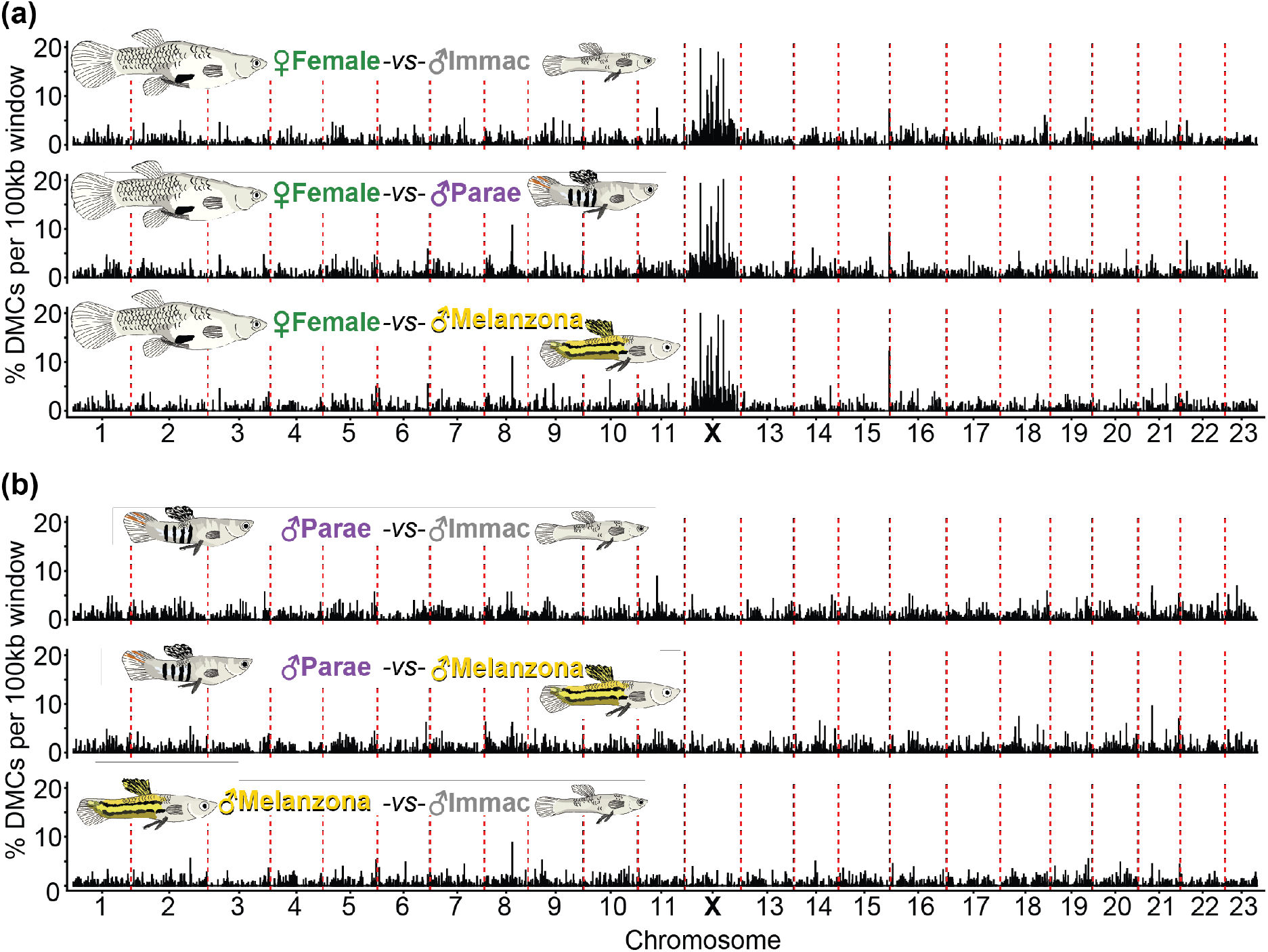
Methylation differences between the sexes are concentrated on the X chromosome, whereas differences among male morphs are spread across the genome. Percentage of CpG sites that are differentially methylated (DMCs) in 100-kb windows along each chromosome for (a) females versus each male morph and (b) each pair of male morphs. Chromosome 12 is the X chromosome. Dashed red lines mark chromosome boundaries. Alt text: Six stacked genome-wide plots, each showing the percentage of differentially methylated CpG sites in 100-kb windows along chromosomes 1–23, with the X chromosome in the position of chromosome 12. The top three plots compare females with each male morph (Immaculata, Parae, Melanzona); in all three, values on the autosomes stay mostly below 5%, while the X chromosome shows a dense block of windows reaching 10–20%. The bottom three plots compare pairs of male morphs (Parae vs. Immaculata, Parae vs. Melanzona, Melanzona vs. Immaculata); values are low and similar across all chromosomes, including the X, with no chromosome standing out. Fish illustrations identify the two groups in each comparison.

### Regional differences: morph and sex DMRs are similarly widespread across autosomes but sex DMRs are more pronounced on the X chromosome

Biologically relevant CpG methylation often correlates spatially among neighboring sites, and regional analysis can reveal coordinated methylation differences that help characterize them relative to nearby genomic features (Affinito et al. 2020). We therefore identified *differentially methylated regions* (DMRs) independently for each sex and male-morph contrast and compared their genomic distribution, size, CpG content, and magnitude.

DMRs showed the same genomic distribution as DMCs, occurring throughout the autosomes in all contrasts and enriched on the X chromosome in sex contrasts (**Supplementary Figures 2, 3**). On the autosomes, sex and male-morph contrasts identified similar numbers of DMRs (3,498–4,121 and 3,971–4,166, respectively). Meanwhile, on the X chromosome, sex contrasts identified roughly twice as many DMRs (314–375) as male-morph contrasts (168–188). DMRs were similar in size and CpG content across all contrasts, averaging 437–460 bp and ∼21 CpGs per region (**Supplementary Table 2**).

The contrasts also differed in the magnitude of methylation differences, and again mainly on the X chromosome. On the autosomes, typical differences were similar between sex and male-morph contrasts (median absolute difference 10.25–12.30% and 11.8–13.06%, respectively). X-linked DMRs in sex contrasts, however, showed larger differences in a consistent direction: across these DMRs, females averaged 69–71% methylation compared with 33–40% in males (median absolute difference 21.36–34.42% and 11.63–13.68%, respectively). This pattern was most pronounced among the strongest DMRs. Each sex contrast contained 115–119 DMRs with a methylation difference greater than 75%, of which 104–110 were on the X chromosome and more highly methylated in females (**Supplementary Figure 2**). In comparison, each male-morph contrast contained a single DMR with a methylation difference greater than 75%, none on the X chromosome (**Supplementary Table 2**).

Thus, male-morph contrasts identified as many autosomal DMRs as sex contrasts, with similar typical magnitudes, but lacked the large, X-linked differences that distinguish the sexes. Because each set of DMRs was identified within a single contrast, we next asked whether the same regions recur across contrasts.

### Phenotype-associated methylation differences recur across contrasts

If a region is differentially methylated due to phenotype rather than individual variation, then every pairwise contrast that includes that phenotype should detect it. For example, a region where Immaculata males differ from all other phenotypes should be detected in the Immaculata–Parae, Immaculata–Melanzona, and female–Immaculata contrasts, but not in the remaining three. Because each phenotype is included in three of the six contrasts, our design allows this prediction to be tested directly.

To apply this test, we merged overlapping DMRs from all six contrasts into consensus regions, hereafter *variably methylated regions* (VMRs), and recorded the combination of pairwise contrasts in which each VMR was differentially methylated (**Figure 2**). Each combination corresponded to a particular grouping of phenotypes: *sex-specific* VMRs, in which females differed from all three male morphs; *morph-specific* VMRs, in which one morph differed from females and from the other two morphs; *sex & morph* VMRs, in which females and one morph each differed from all other phenotypes; and *cross-phenotype* VMRs, in which two phenotypes share a methylation state that differed from the other two. We interpret the VMRs detected in only one contrast (*pairwise-specific*), and remaining combinations that do not correspond to a consistent grouping of phenotypes (*other*) as more likely to reflect individual variation or detection threshold noise.

**Figure 2.**
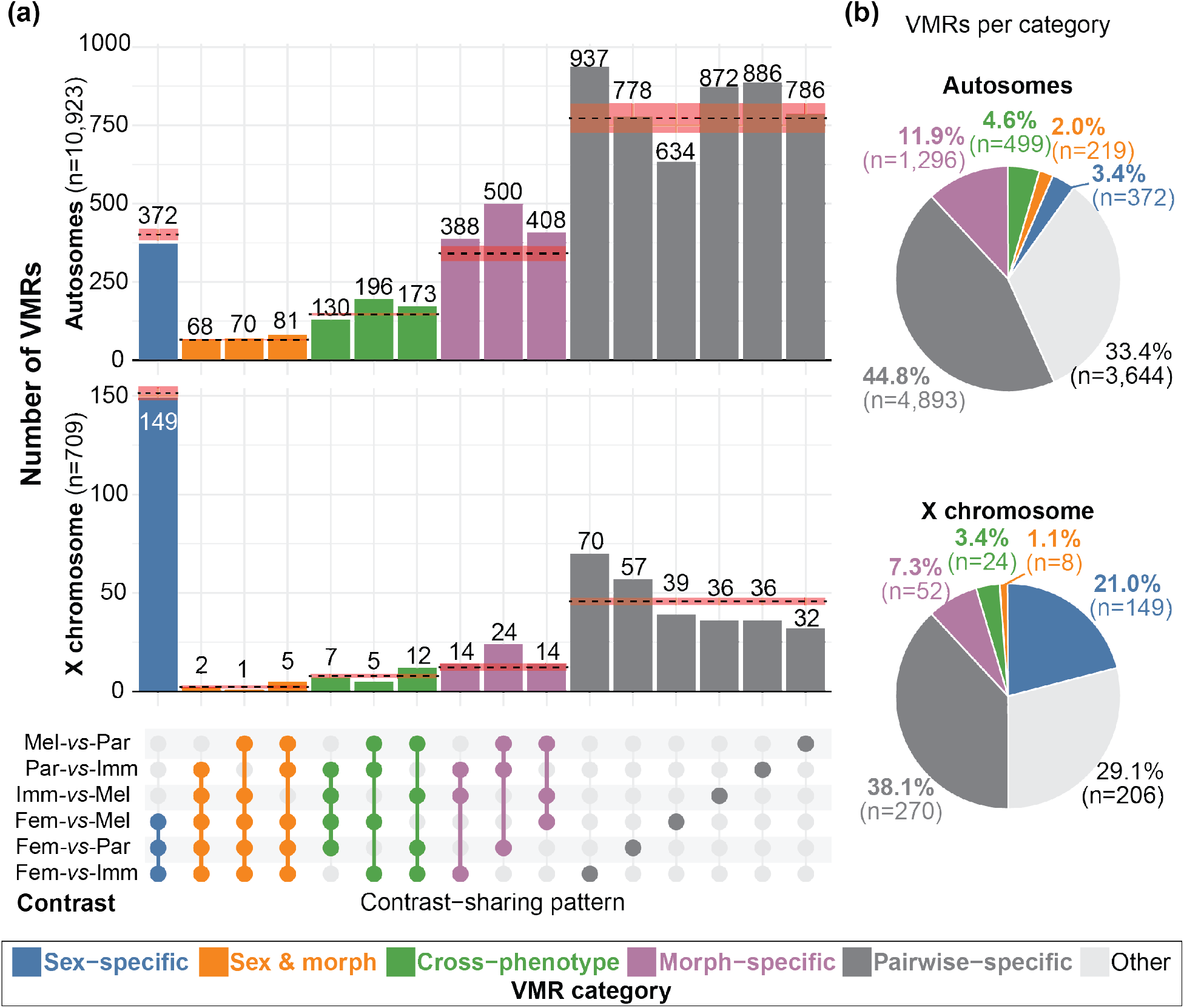
Morph-specific methylation differences recur across comparisons more often than expected by chance. (a) Number of variably methylated regions (VMRs; merged DMRs) detected by each combination of the six pairwise comparisons (dot matrix), on the autosomes (top) and X chromosome (bottom). Colors indicate VMR categories (defined in the text). Dashed lines and red shading show the mean ± 1 SD expected by chance, from the same six comparisons repeated with males randomly assigned to pseudo-morphs containing one male of each morph (Supplementary Figure 4, Supplementary Table 3); within a category, all combinations share one expected value. (b) Proportion of VMRs in each category, including combinations not shown in (a) (Other). Alt text: Panel a is an upset plot, showing how many variably methylated regions were detected by each combination of the six pairwise comparisons, separately for autosomes (top, 10,923 regions) and the X chromosome (bottom, 709 regions). Bars are colored by category: sex-specific, sex and morph, cross-phenotype, morph-specific, and pairwise-specific. Dashed lines with shaded bands mark the number expected from randomized groupings. On the autosomes, the three morph-specific bars (388, 500, and 408) all exceed the expected value of roughly 340, whereas sex-specific regions (372) match the randomized expectation. On the X chromosome, the sex-specific bar (149) is far taller than any morph-specific bar (14–24). Panel b shows two pie charts of the share of regions in each category: pairwise-specific regions are the largest share on both the autosomes (44.8%) and the X (38.1%), and sex-specific regions make up 21.0% of X-linked but only 3.4% of autosomal regions.

Contrasts involving the same phenotype are not independent, because they share the same three individuals. With only three individuals per group, a single individual with unusually high or low methylation at a region could cause that region to be called a DMR in all three contrasts involving that phenotype. To estimate how often recurrence arises this way, we created *pseudo-morphs*: groups of three males containing one male of each morph. We divided the nine males into three such pseudo-morphs in nine different ways, giving nine balanced shuffled sets (27 pseudo-morphs in total; **Supplementary Table 3, Supplementary Figure 4**). Within each set, we repeated the original design, calling DMRs between each pair of pseudo-morphs and between each pseudo-morph and females, and classified the resulting VMRs into the same categories as the real data. Because every pseudo-morph contains all three morphs, any recurrence across contrasts sharing a pseudo-morph reflects individual variation rather than morph identity. For each category (sex-specific, morph-specific, etc.), we calculated enrichment as the number of VMRs in the real groups divided by the mean number across the nine shuffled sets.

On the autosomes, the contrasts between true morph groups yielded 10,923 VMRs, compared with 9,641–10,937 in the randomized pseudo-morph contrasts. Pairwise-specific VMRs were the most common category in both datasets (44.8% and 43.7–47.4 %), consistent with substantial individual variation within each contrast. Morph-specific VMRs were ∼1.3-fold more frequent in the real data (n = 1,296; 11.9% of autosomal VMRs) than among pseudo-morphs (n = 913-1112). Because female identity was preserved in the randomization, sex-specific VMRs were expected to persist in the randomized data. They did, at similar numbers (372 real vs. 364–429 randomized), confirming that the randomization removed morph structure but not sex structure.

On the X chromosome, the contrasts between true morph groups yielded 709 VMRs (635–690 randomized), of which 38.1% were pairwise-specific (39.4–43.0% randomized). Morph-specific VMRs (n = 52; 7.3%) were ∼1.4-fold more frequent than among pseudo-morphs (n = 29–46), similar to the enrichment on the autosomes. Sex-specific VMRs, however, made up a far larger share of X-linked VMRs (n = 149; 21.0%) than of autosomal VMRs (3.4%), and outnumbered each morph-specific pattern (Immaculata, 14; Melanzona, 14; Parae, 24) by 6–11 fold.

Together, these results show that methylation differences recur in morph-specific patterns more often than expected from individual variation alone, on both the autosomes and the X chromosome. In contrast, sex-specific differences are disproportionately concentrated on the X chromosome.

### Morph-specific and sex-specific methylation patterns differ between the autosomes and X chromosome

To examine the methylation states underlying these recurrent patterns, we plotted the methylation level of each individual at sex-specific and morph-specific VMRs (**Figure 3**). Because these VMRs were already identified from phenotype contrasts, individuals are expected to group by phenotype. The heatmaps therefore illustrate how phenotypes differ rather than test whether they do. At both autosomal and X-linked VMRs, the three individuals of each phenotype showed largely consistent methylation states, indicating that the recurrent patterns were not driven by single individuals.

**Figure 3.**
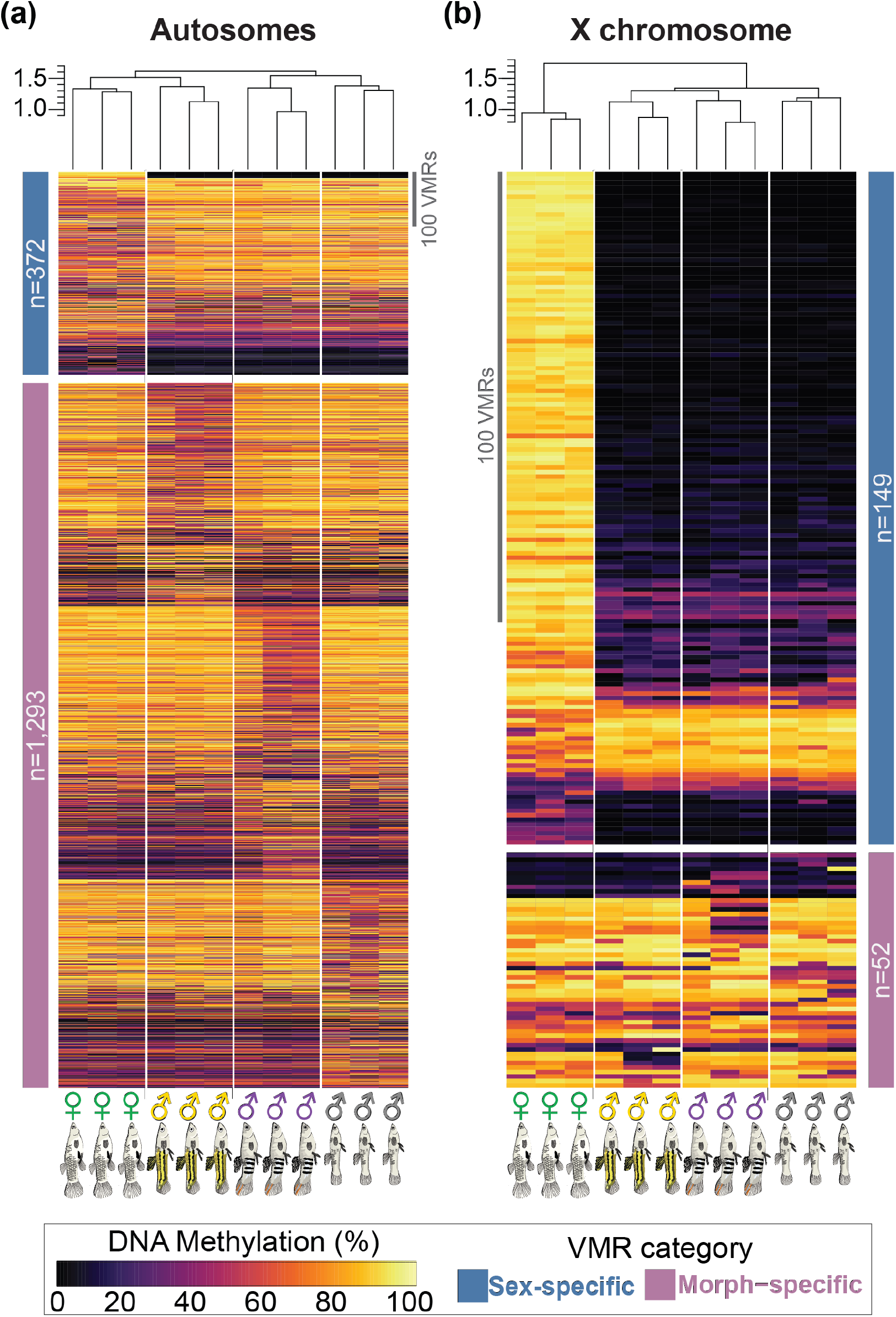
Individuals of the same phenotype show consistent methylation at sex-specific and morph-specific VMRs. Methylation level of each individual (columns) at each sex-specific (blue) and morph-specific (pink) VMR (rows; categories as in Figure 2; VMRs with missing data or <10% difference across individuals are not shown) on (a) the autosomes and (b) the X chromosome. Fish symbols indicate females (green) and Melanzona (yellow), Parae (purple), and Immaculata (grey) males. Rows are ordered by the phenotype that most distinguishes each VMR (see Methods). Dendrograms show clustering of individuals based on all VMRs in each genomic region (see Methods). Alt text: Two heatmaps of methylation level (0–100%, black to yellow) in 12 individuals (columns: three females, then three each of Melanzona, Parae, and Immaculata males) at sex-specific and morph-specific variably methylated regions (rows). Panel a shows the autosomes (372 sex-specific and 1,293 morph-specific regions); panel b shows the X chromosome (149 sex-specific and 52 morph-specific regions). A dendrogram of each heatmap groups individuals of the same phenotype together, and across rows the three individuals show similar colors, producing blocks of consistent methylation. On the X chromosome, most sex-specific regions are highly methylated in all three females and nearly unmethylated in all nine males. On the autosomes, sex-specific and morph-specific regions include both higher and lower methylation in the distinguishing phenotype.

The methylation direction of these differences, however, differed between the autosomes and the X chromosome. On the autosomes, sex-specific and morph-specific VMRs included both higher and lower methylation in the distinguishing phenotype, and individual VMRs spanned a broad range of methylation levels (**Figure 3A**). On the X chromosome, by contrast, most sex-specific VMRs (104 of 149) were highly methylated in females and nearly unmethylated in all three male morphs, whereas X-linked morph-specific VMRs resembled their autosomal counterparts (**Figure 3B**). In both categories, VMRs were distributed along all chromosomes rather than concentrated in a few regions (**Supplementary Figure 5**).

### VMRs overlap with genes more often than expected by chance

To investigate the genomic context of VMRs, we assessed their positions relative to genes. More VMRs overlapped genes than expected from the proportion of the genome occupied by genes, both genome-wide (7,118 observed vs. 6,755 expected; pooled *X^2^* = 46.52, df = 1, p = 9.12 x 10^-12^) and on the autosomes (6,731 vs. 6,375; pooled *X^2^* = 47.52, df = 1, p = 5.43 x 10^-12^). This excess was not driven by a few chromosomes: deviations from expectation remained significant when expected counts were calculated separately for each chromosome (chromosome-aware *X^2^*= 102.76, df = 23, p = 4.7 x 10⁻¹² genome-wide; *X^2^* = 100.95, df = 22, p = 4.4 x 10⁻¹² for autosomes) (**Supplementary Figure 6**). This suggests VMRs are preferentially located in genes, where they could affect gene regulation.

### Gene methylation is consistently but weakly associated with expression

Given the spatial association between gene bodies and VMRs, we next examined how methylation relates to gene expression. For this analysis, we used 3’ RNA-seq data from tail muscle of different individuals from the same laboratory population (three per phenotype; see Methods). In vertebrates, promoter methylation is generally associated with transcriptional repression (Fradin et al. 1982; Jones and Takai 2001; Maunakea et al. 2010; Shukla et al. 2011), whereas gene body methylation often shows an inverted-U relationship with expression, with the highest expression at intermediate methylation (Feng et al. 2010; Zemach et al. 2010). Within each phenotype, we indeed found the same inverted-U-shaped relationship between gene expression and DNA methylation across the gene **(Supplementary Figure 7).**

We then asked whether methylation differences between phenotypes were associated with expression differences. For each gene and contrast, we regressed the absolute log₂ fold change in expression on the absolute difference in gene-body methylation (**Supplementary Figure 8**). The relationship was positive in all six autosomal contrasts (FDR < 0.0001) and in five of six X-chromosome contrasts, all except Melanzona vs. Parae (**Supplementary Table 4**). Because the X chromosome contains far fewer genes than the autosomes (595 vs. ∼14,140), and in the three sex contrasts its genes also showed larger methylation differences, we tested whether either factor shaped the autosomal result. We repeatedly subsampled autosomal genes (5,000 times per contrast), first to the number of X-linked genes, and then to both their number and their distribution of methylation differences. The autosomal slope remained positive in 97.6–99.4% and 98.0–99.8% of subsamples, respectively, indicating that the direction of the relationship did not depend on gene number or on the range of methylation differences.

These relationships were consistent but weak. Independent linear regressions for each contrast and genomic partition showed that methylation differences in the gene body explained at most 1.8% of the variation in expression differences on the autosomes (R^2^ = 0.018) and up to 8.5% (R^2^ = 0.085) on the X chromosome. Slopes generally did not differ between the autosomes and the X chromosome; the methylation-by-chromosome interaction was significant after FDR correction only for Parae vs. Immaculata, where the relationship was stronger on the X (**Supplementary Table 4**). Thus, methylation differences track expression differences across the genome but account for only a small proportion of expression divergence.

## DISCUSSION

At the whole-genome scale, we found that both sex and alternative reproductive morphs are associated with distinct DNA methylation landscapes in *P. parae*. These results are consistent with the long-standing theoretical expectation that alternative epigenetic states in the shared genome may contribute to the coordination of complex alternative phenotypes (West-Eberhard 1986; Oliveira et al. 2008).

### Morph and sex define distinct axes of methylome divergence

Methylation differentiation among *P. parae* morphs was broadly distributed across the autosomes, with numerous regional differences varying in location and magnitude rather than concentrated as a uniform shift toward higher or lower methylation. This broadly distributed pattern is consistent with the complexity of alternative male morphs, which strongly differ in morphology, reproductive strategy, and behavior (Sandkam 2026). As originally proposed by West-Eberhard (1986) alternative phenotypes arise when a regulatory “switch” channels a shared genome into discrete developmental and phenotypic trajectories, coordinating suites of otherwise independent traits. Such channeling does not require a single, uniform directional change across the genome. Consequently, the observed distribution of morph-associated methylation differences across the shared genome is compatible with the complex regulatory programs required by alternative phenotypes.

Previous studies have linked genome-wide DNA methylation differences to alternative life-history strategies and phenotypic diversification in multiple vertebrate systems (Venney et al. 2023). However, among genetically determined polymorphisms, the genomic distribution of morph-associated differentiation does not seem to follow a single pattern. In white-throated sparrows (*Zonotrichia albicollis*) for example, methylation differences between alternative plumage and behavioral morphs are predominantly concentrated within the nonrecombining chromosomal region underlying morph identity (Sun et al. 2021; Prichard et al. 2022). In comparison, in *P. parae*, where male morph identity is also linked to alternative Y-chromosome haplotypes (Lindholm et al. 2004; Sandkam et al. 2021), we found broad morph-associated methylation differences extending across the shared autosomal genome. This contrast suggests that genetically determined alternative phenotypes can exhibit distinct spatial architectures of methylation differentiation.

Unlike the morph pattern, sex-associated methylation differences in *P. parae* follow a more expected organization: females are more highly methylated than males across much of the X chromosome. This structure is consistent with previous studies linking sex-chromosome evolution and dosage-related processes to chromosome-wide methylation differentiation in poeciliids (Darolti et al. 2019; Metzger et al. 2021; Metzger et al. 2023; Fong et al. 2025; Metzger and Mank 2026). Recovering this pronounced female-male axis provided an internal biological benchmark for our analyses and emphasizes that the morph-associated autosomal pattern represents a qualitatively different form of methylome divergence. Together, these patterns show that a single genome can carry two distinct forms of methylome divergence: broadly distributed autosomal differences among male morphs, comparable in number to those between the sexes, and an additional chromosome-wide difference between the sexes on the X.

### Alternative male methylomes arise under common-garden conditions

Environmental effects on methylation are well documented. In *Astatotilapia burtoni*, experimentally altered visual environments modify methylation and expression of the endothelin receptor B pathway (Fang et al. 2025), while freshwater and marine three-spined stickleback exhibit divergent methylation landscapes containing both heritable and salinity-responsive components (Smith et al. 2015; Artemov et al. 2017; Metzger and Schulte 2018; Heckwolf et al. 2020; Hu et al. 2021). In such systems, environmental differences often overlap with phenotype-associated methylation, complicating the identification of the heritable epigenetic variation associated with alternative phenotypes. This potential confounding effect is minimized in this study with *P. parae*, as all individuals originated from the same laboratory population and were maintained in the same recirculating water system, under common-garden conditions. Because individuals of each morph were drawn at random from stocks, the recurrent morph-specific differences we observe are unlikely to reflect environmental variation. Instead, the reported methylation profiles reflect reproducible alternative methylation landscapes associated with inherited male morph identity. These profiles support DNA methylation as an additional layer of molecular differentiation accompanying the alternative male phenotypes of *P. parae*.

### Methylation and gene expression

In vertebrates, DNA methylation in gene-coding sequence has been associated with transcriptional activity, chromatin state, alternative promoter function, and alternative splicing (Lister et al. 2009; Zemach et al. 2010), and the strength and direction of the relationship between methylation status and gene expression vary across tissues, genomic contexts, and taxa (Lister et al. 2009; Metzger et al. 2023; Ross et al. 2023; Metzger and Mank 2026). In *P. parae*, gene methylation within each phenotype showed an inverted-U relationship with expression, with the highest expression at intermediate methylation, consistent with patterns reported for gene-body methylation in other vertebrates (Feng et al. 2010; Zemach et al. 2010). Differences in gene methylation between phenotypes were also consistently, though weakly, associated with differences in expression, suggesting that morph-associated methylation differences may contribute modestly to transcriptional divergence.

The weakness of this association is not unexpected. Transcriptional regulation reflects multiple interacting layers of epigenetic control (Cedar et al. 2022), and in other systems, broad methylation differences between populations or lineages often occur without correspondingly large differences in gene expression (Venney et al. 2023). In addition, methylation and expression were measured in different individuals, so our comparison relies on phenotype means and likely underestimates within-individual association. Morph-associated methylation differences may therefore contribute modestly or indirectly to transcriptional divergence rather than driving it alone. Future studies should combine matched methylome and transcriptome data from the same individuals to test the causal relationships behind these differences. This would be especially informative across the developmental stages and tissues in which alternative male phenotypes are established, because morph differences likely emerge through coordinated epigenetic programs acting across time, tissue, and genomic context.

### Conclusions

Our results show that inherited male morphs in *P. parae* are accompanied by alternative DNA methylation landscapes in adult somatic tissue. Morph-associated methylation divergence is distributed across the shared autosomal genome and includes morph-specific regions that recur more often than expected from individual variation. Sex-associated methylation divergence follows a comparable pattern on the autosomes but forms an additional, chromosome-wide axis on the X chromosome, consistent with a role for methylation in X-chromosome regulation. Methylomes have often been compared across species, populations, environments, developmental stages, tissues, and cell types (Eckhardt et al. 2006; Lister et al. 2009; Bogdanović et al. 2016; Metzger and Schulte 2018; Heckwolf et al. 2020); the morphs of *P. parae* show that multiple discrete methylome configurations can also coexist within a single species, sex, tissue, and adult developmental stage. Many systems with discrete phenotypes have well-characterized genetic bases, including sex-limited polymorphisms, supergene-associated morphs, and alternative reproductive tactics that are associated with broader differences in genome usage (Oliveira et al. 2008; Mank 2023). *P. parae* offers a natural model for investigating how such inherited polymorphisms are accompanied by genome-wide epigenetic divergence.

## MATERIALS AND METHODS

### Fish Maintenance and DNA extraction

We used individuals from an outbred laboratory colony of *Poecilia parae*, originally collected from Suriname and maintained in Ithaca, NY. Fish were reared in large communal tanks, separated by morph, under standardized conditions: a constant temperature of 24 °C, a 12:12 h light cycle, continuous recirculating water flow, and *ad libitum* feeding with a combination of commercial fish food pellets (Fancy Guppy – Hikari) and live brine shrimp, provided twice daily.

Three adult individuals were sampled from each of the three male major-morphs (Immaculata, Parae, and Melanzona) and females (n=12 total). All fish were euthanized by overdose of MS-222 diluted in system water (3.33 mg/mL) (Cornell IACUC protocol #2021-0067). Tail muscle tissue was dissected in phosphate-buffered saline (PBS), with the tissue sample taken from the region spanning the posterior edge of the anal pore to the base of the caudal fin. Genomic DNA was extracted using the Monarch Genomic DNA Purification Kit (NEB #T3010).

### Library Prep and Sequencing

To detect methylated cytosines in our samples, we generated individual libraries using ≥200 ng of genomic DNA per sample and spiked in two internal controls to verify enzymatic conversion: unmethylated lambda DNA and CpG-methylated pUC19. Libraries were constructed using NEBNext Enzymatic Methyl-seq (EM-seq) (NEB #E7120S) according to manufacturer’s instructions (Vaisvila et al. 2021). All libraries were sequenced together on both an Illumina NextSeq 2000 (2×100 bp PE reads) and on a NovaSeqX platform (2×150bp PE reads) by the Cornell Genomics Core Facility.

### Genome Alignment and Cytosine Methylation Calling

Pooled reads from both sequencing runs were aligned to the female *Poecilia parae* reference genome (chromosome 12 is the X chromosome). To assess enzymatic conversion efficiency and overall library quality, we added the lambda and pUC-19 sequences as independent scaffolds to the reference genome. We used the nf-core/methylseq pipeline (v4.2.0) with the ‘--em_seq’ flag (Di Tommaso et al. 2017; Ewels et al. 2020; Ewels et al. 2024). This pipeline merges FASTQ files from the two sequencing runs for each sample, trims reads with Trim Galore (v0.6.7), assesses read quality with FastQC (v0.12.1), and generates alignments using Bismark (v0.24.0) (Krueger and Andrews 2011). Reads were deduplicated with ‘deduplicate_bismark’. It provided a per-sample cytosine report for all cytosines in the genome, including their genomic positions, sequence contexts, coverage, and methylation frequencies.

We achieved a mean coverage of 20.9X **(Supplementary Figure 1A),** which is typical of comparable whole-genome studies investigating alternative methylation patterns within species (Wang et al. 2015; Bain et al. 2021; Pan et al. 2021; Singh et al. 2021; Yang et al. 2021). Of all sequenced cytosines, 9.3% were in the CpG context, yet methylation was largely restricted to this context: CpG sites were 74.46 ± 0.26% (mean ± SEM) methylated, compared with 0.53% and 0.52% in CHG and CHH contexts (**Supplementary Figure 1B,C**). This pattern is consistent with cytosine-context distributions reported in vertebrates and other teleost fishes (Duncan et al. 2018; Metzger and Schulte 2018; Matlosz et al. 2022; Zhou et al. 2022; Klughammer et al. 2023; Metzger et al. 2023), in which methylation occurs predominantly at CpG sites (Bird 1986; Feng et al. 2010). We therefore restricted all subsequent analyses to CpGs. As CpG loci are symmetrical, we collapsed counts from both strands into a single strand, using the ‘sym’ function of DNMTools (v1.2.2; Song et al., 2013).

Spike-in controls confirmed efficient and specific conversion: unmethylated DNA in the lambda control was converted at >99.99% efficiency, whereas ≤1.16% of methylated CpGs in the pUC19 control were converted **(Supplementary Figure 1D).** As with all conversion-based methods, C/T polymorphisms at CpG sites cannot be distinguished from unmethylated cytosines; however, allele frequencies were not expected to differ between sexes/morphs and were therefore unlikely to affect differential methylation.

### Genome-Wide Methylation Structure

To verify whether genome-wide CpG methylation profiles were associated with phenotype identity, we analyzed CpG methylation levels from the 12 individuals. CpGs were initially required to have a minimum sequencing coverage (≥5 reads) and only sites with methylation values in all 12 samples were retained for the analyses (15,806,357 total; 11,323,952 retained). Pairwise dissimilarity between samples was calculated as the mean absolute difference in methylation level across all retained CpGs, such that smaller values indicated more similar methylation profiles.

Phenotype-associated differences in multivariate methylation profiles were tested using PERMANOVA, with phenotype as a four-level factor and significance assessed by permuting phenotype labels (9,999 permutations). To determine whether a significant PERMANOVA result could be influenced by differences in within-group variability, homogeneity of multivariate dispersion among phenotype groups was evaluated using PERMDISP with 9,999 permutations.

Additionally, we directly tested whether individuals of the same phenotype had more similar methylation profiles than individuals of different phenotypes. To do so, we calculated the difference between the mean pairwise distance among individuals of different phenotypes and the mean pairwise distance among individuals of the same phenotype; a positive difference indicates that individuals of the same phenotype are more similar. We assessed significance by comparing this observed difference with the differences obtained after randomly permuting phenotype labels among individuals 9,999 times (one-sided test). PERMANOVA and PERMDISP were run with the ‘vegan’ package (Oksanen et al. 2026), and the distance test was implemented in base R.

### Detection of Differential Methylation

To identify *differentially methylated cytosines* (DMCs), we used the *radmeth* function in DNMTools (v1.2.2; (Dolzhenko and Smith 2014), which fits a beta-binomial regression to methylated and total read counts, with phenotype as the factor in each pairwise contrast. P-values were then combined with those of neighboring CpGs within 200bp using *radadjust* (a weighted Z-test) and corrected for multiple testing; CpGs with a corrected p-value ≤ 0.01 were considered DMCs. To visualize the genomic distribution of DMCs **(Figure 1)**, we calculated the percentage of sequenced CpGs called as DMCs in 100-kb windows, retaining windows with at least 10 sequenced CpGs. DMC counts were summarized separately for the whole genome, the autosomes, and the X chromosome **(Supplementary Table 1)**.

To identify precise differentially methylated regions (DMRs), we used Methylasso (v 1.0.0)(Balaramane et al. 2024) with the parameters: minimum coverage of 5 reads (– c), minimum methylation difference of 10% (–d 0.1), DMR significance threshold < 0.0001 (–p), maximum distance for merging segments (500bp), minimum width for segments (100bp), and default settings otherwise (≥4 CpGs per DMR).

To characterize the degree of differential methylation between phenotypes in each contrast, we used the methylation difference of each DMR calculated with Methylasso to plot histograms of methylation differences between the groups. We visualized skew in the genomic distribution of DMRs and their methylation differences in each contrast using scatter plots of methylation level per DMR. All plots were generated using ‘ggplot2’ in R (Wickham 2016).

### Variably Methylated Regions

To compare differentially methylated regions across pairwise contrasts, we combined all DMRs from the contrasts and merged those that overlapped or touched (bedtools merge –d 0; Quinlan & Hall, 2010), hereafter referred to as *variably methylated regions* (VMRs). A DMR found in only one contrast formed a VMR on its own. For each VMR, every contrast was then scored as to whether there was a DMR overlapping with it and compared between all six contrasts using the ‘complexupset’ package in R (Krassowski et al. 2022). To compare methylation levels of VMRs between samples, we calculated the mean methylation of all CpGs within the VMR covered by at least one read using ‘bedtools map’ v2.30.0 (Quinlan and Hall 2010).

### Sample-Identity Randomization Tests

To estimate how often contrast-sharing patterns arise by chance, we repeated the DMR and VMR analyses after assigning the nine males to three pseudo-morphs of three individuals, each containing one male of each morph (27 pseudo-morphs, 3 pseudo-morphs per set, 9 sets in total) **(Supplementary Figure 4).** For each set, we ran the same six contrasts as in the real design (each pseudo-morph against the other two and against females) with identical parameters, and merged the resulting DMRs into a separate set of pseudo-VMRs, so that VMR boundaries in real and randomized data were defined in the same way **(Supplementary Figure 4)**. Because female identity was preserved, these contrasts retain sex differences but not morph differences. For each contrast-sharing category, enrichment was calculated as the number of VMRs in the real data divided by the mean number of pseudo-VMRs across the nine randomized sets.

### Heatmaps of VMR Methylation

Heatmaps were generated separately for the autosomes and X-chromosome using pheatmap (Kolde 2025), with data.table (Barrett et al. 2026) for data manipulation and viridis (Garnier et al. 2024) for the methylation color scale in R. VMRs with any missing samples or whose methylation range across all 12 fish was under 10% were removed. Hierarchical clustering was applied only to samples (heatmap columns). Each VMR was standardized across samples using Z-scores, and pairwise sample distances were calculated as Euclidean distances across the full set of standardized VMR profiles within each genomic partition, divided by the square root of the number of VMRs. Samples were then hierarchically clustered using complete linkage, and the resulting dendrogram was retained as a fixed sample tree. The heatmap tile colors preserve the original, unscaled methylation values for interpretability.

VMR rows were organized by biological contrast-sharing categories derived from the Upset plots, rather than by hierarchical clustering. Within each category, phenotype associations were quantified using the group-equalized point-biserial correlation (r.g) implemented in the ‘indicspecies’ R package (Tichy and Chytry 2006; Cáceres and Legendre 2009; De Cáceres et al. 2012) across Z-scores. For each VMR, ‘strassoc’ was used to calculate its association with Female, Immaculata, Parae, and Melanzona phenotypes; the phenotype with the strongest absolute association was assigned as the indicator phenotype. VMRs were then grouped by indicator phenotype and ordered by decreasing signed r.g within each group. These association statistics were used only to organize rows and not to filter VMRs. Additional summaries of the genomic locations of all VMRs were generated using ggplot2 (Wickham 2016).

### Genomic Association of VMRs with Genes

To characterize the spatial relationship between VMRs and annotated genes, we compared their locations in the *P. parae* genome. For each VMR, the closest annotated gene was identified with ‘*bedtools closest’* (–D b –t all) using strand-aware signed distances, such that overlapping VMRs were assigned distance 0, upstream regions negative distances, and downstream regions positive distances (Quinlan and Hall 2010). VMRs were then classified as intragenic or intergenic.

To test whether VMR occupancy in genes matched the genomic availability of genic sequence, we compared the observed VMR overlap with gene models against the expectations derived from gene occupancy of the genome alone. We tested this with both a pooled X^2^ test and with a chromosome-aware X^2^ test, in which expected intragenic and intergenic counts were calculated separately for each chromosome from its gene occupancy and X^2^ deviations were summed across chromosomes. Both tests were run for the whole genome and for the autosomes alone.

### Relation of DNA Methylation and Gene Expression

To evaluate the relation between DNA methylation and gene expression we compared the DNA methylation level to the gene-level expression counts from adult muscle tissue generated as part of another study on this same lab population (control (untreated) individuals from Taves et al – *in revision*) (3’ RNA-seq reads available on NCBI: Bioproject PRJNA1415236 – providing three samples per phenotype).

Raw 3’ RNA-seq reads were processed using ‘bbduk.sh’ (BBTools v39.08) (Bushnell 2014). Adapter sequences were removed using ‘forcetrimleft=12, ktrim=r, k=13, and mink=5’, followed by quality trimming and filtering using ‘qtrim=r, trimq=20, minlen=50, trimpolya=5, and trimpolyg=5’. Read quality and adapter content were assessed before and after trimming using FastQC (v0.12.1) (FastQC 2015). Retained reads were aligned to the annotated *P. parae* female genome using STAR (v2.7.11b) (Dobin et al. 2013). Alignments were restricted to uniquely mapping reads, with a minimum matched fraction of 33% of the read length and a maximum of one reported alignment per read. Gene-level counts were obtained directly during alignment using ‘–– quantMode GeneCounts’, retaining counts corresponding to the forward-strand library orientation. After filtering and alignment, a mean of approximately 6.7 million uniquely mapped reads was retained per sample.

For quality control of gene-expression profiles, count matrices were variance-stabilized using ‘varianceStabilizingTransformation’ in DESeq2 (v1.46.0) (Love et al. 2014), and potential sample outliers were evaluated using ‘arrayQualityMetrics’ (v3.62.0). Sample clustering was additionally assessed using Poisson distances calculated with ‘PoiClaClù (v1.0.2.1) (Witten 2019) in R.

Gene expression counts were normalized to Counts per Million (CPM) to account for library size differences. To avoid undefined values at zeros and stabilize variance across expression ranges, values were transformed as log2(CPM+1). For each sample, we calculated the methylation level across the full gene body. We defined the gene body as the genomic sequence within the annotated gene boundaries, excluding the 5′ and 3′ untranslated regions (UTRs). This definition encompasses both protein-coding sequences (CDS) and intronic regions.

Associations between methylation and expression on autosomes and the X chromosome were evaluated independently for each morph and females. Linear relationships were summarized per gene, by ordinary least-squares regression of the mean log₂(CPM+1) across the three individuals of a phenotype against that phenotype’s mean gene-body methylation, with model fit reported as R^2^. Monotonic associations were assessed using Spearman rank correlation (cor.test, method = “spearman”, exact = FALSE) (Kassambara 2023). To test for non-linear unimodal relationships, we fit quadratic models of the form y=β_0_+(β_1_*x)+(β_2_*x^2^). Evidence for an inverted-U pattern was inferred from a negative quadratic coefficient, support for the quadratic term, and an improved fit relative to the linear model (ANOVA). As a flexible non-linear diagnostic, generalized additive models were also fit with ‘mgcv’ (Wood 2017) using s(x, k = 5) and REML estimation.

### Differential Methylation and Differential Expression Analysis

Differential gene expression analyses were performed using the DESeq2 package (v1.46.0) in R (Love et al. 2014). Before fitting, genes were filtered using a global CPM threshold implemented with edgeR (CPM ≥ 1 in at least 3 samples) (Robinson et al. 2010), followed by a pair-specific per-group filter requiring CPM ≥ 1 in at least 2 samples per group. DESeq2 Wald tests were used to estimate gene-wise effects, with significance summarized at FDR = 0.05. For effect-size interpretation, raw log_2_ fold changes were additionally shrunk with lfcShrink(…, type = “apeglm”), and shrunken log_2_ fold changes were used in downstream methylation–expression analyses. Gene-by-gene p-values were corrected for multiple testing using Benjamini-Hochberg FDR.

To test whether the relationship between methylation differentiation and gene-expression divergence differed between autosomes and the X chromosome, we used genes with complete methylation and gene-expression information. For each gene, we calculated methylation differentiation as the absolute difference in mean methylation between the two compared phenotypes, and expression divergence as the absolute log_2_ fold change. Chromosome-specific relationships were first evaluated using linear models containing absolute difference in mean methylation, chromosome class, and their interaction (absolute log_2_ fold change ∼ absolute difference in mean methylation x chromosome). The interaction term provided a direct test of whether regression slopes differed between autosomes and the X chromosome.

To determine whether autosomal and X-linked genes differed in the magnitude of methylation differentiation between phenotypes, we compared their distributions of absolute differences in mean methylation using two-sample Kolmogorov–Smirnov and Wilcoxon rank-sum tests. We then asked whether statistical support for the autosomal methylation–expression relationship could be explained by the larger number of autosomal genes. Hence, we randomly downsampled autosomal genes without replacement to the number of X-linked genes and repeated the regression 5,000 times for each contrast. To further assess whether this relationship could be influenced by differences in the magnitude of methylation differentiation between chromosome classes, we performed an additional 5,000 resampling iterations in which autosomal genes were matched to X-linked genes in both number and distribution of methylation differentiation using quantile-based strata. We used the resulting distributions of slopes, R² values, and p-values to evaluate whether the autosomal relationship persisted under both conditions. Benjamini–Hochberg false-discovery-rate correction was applied separately across contrasts for each statistical test.

### AI Usage Disclosure

ChatGPT (OpenAI) and Claude (Anthropic) were used to help write and debug analysis scripts (e.g., R plotting and data-handling code). Claude (Anthropic) was used to edit manuscript text for clarity and grammar, and to cross-check that analyses described in the manuscript matched the code in the project repository. The authors tested all AI-assisted code, and the authors reviewed, revised, and approved all AI-suggested text. No AI tool was used to generate or interpret data.

## DATA AVAILABILITY

EM-seq and RNA-seq reads are available on the Sequence Read Archive (SRA) under NCBI Bioprojects: <u>PRJNA1533486</u> and <u>PRJNA1415236,</u> respectively. All scripts used in this project are available in the GitHub repository (https://github.com/mz448/Poecilia-parae_DNA-methylation.git). Original illustrations of *Poecilia parae* male morphs and females used in this study are publicly available through Figshare (DOI: 10.6084/m9.figshare.28478429) under the Creative Commons Attribution 4.0 International license.

## ACKNOWLEDGEMENTS

We thank Erika Mudrak of the Cornell Statistical Consulting Unit in the Ann S. Bowers College of Computing and Information Science for her thoughtful advice on the statistical analysis. We also thank Carolina Gomez, Helen Stec, and members of the Sandkam lab for the detailed revisions of the initial manuscript. We thank the Genomics Facility (RRID: SCR_021727) of the Biotechnology Resource Center at the Cornell Institute of Biotechnology for assistance with sequencing experiments. This work was supported by the National Institute of General Medical Sciences of the National Institutes of Health under Award Number R35GM155164. The content is solely the responsibility of the authors and does not necessarily represent the official views of the National Institutes of Health. We used ChatGPT (OpenAI) and Claude (Anthropic) to assist with code development and text editing, as described in the Materials and Methods; the authors take full responsibility for the content of this article.

