## Supplementary Figure for "Alternative reproductive morphs and sexes differ in DNA methylation along distinct genomic axes in *Poecilia parae*"

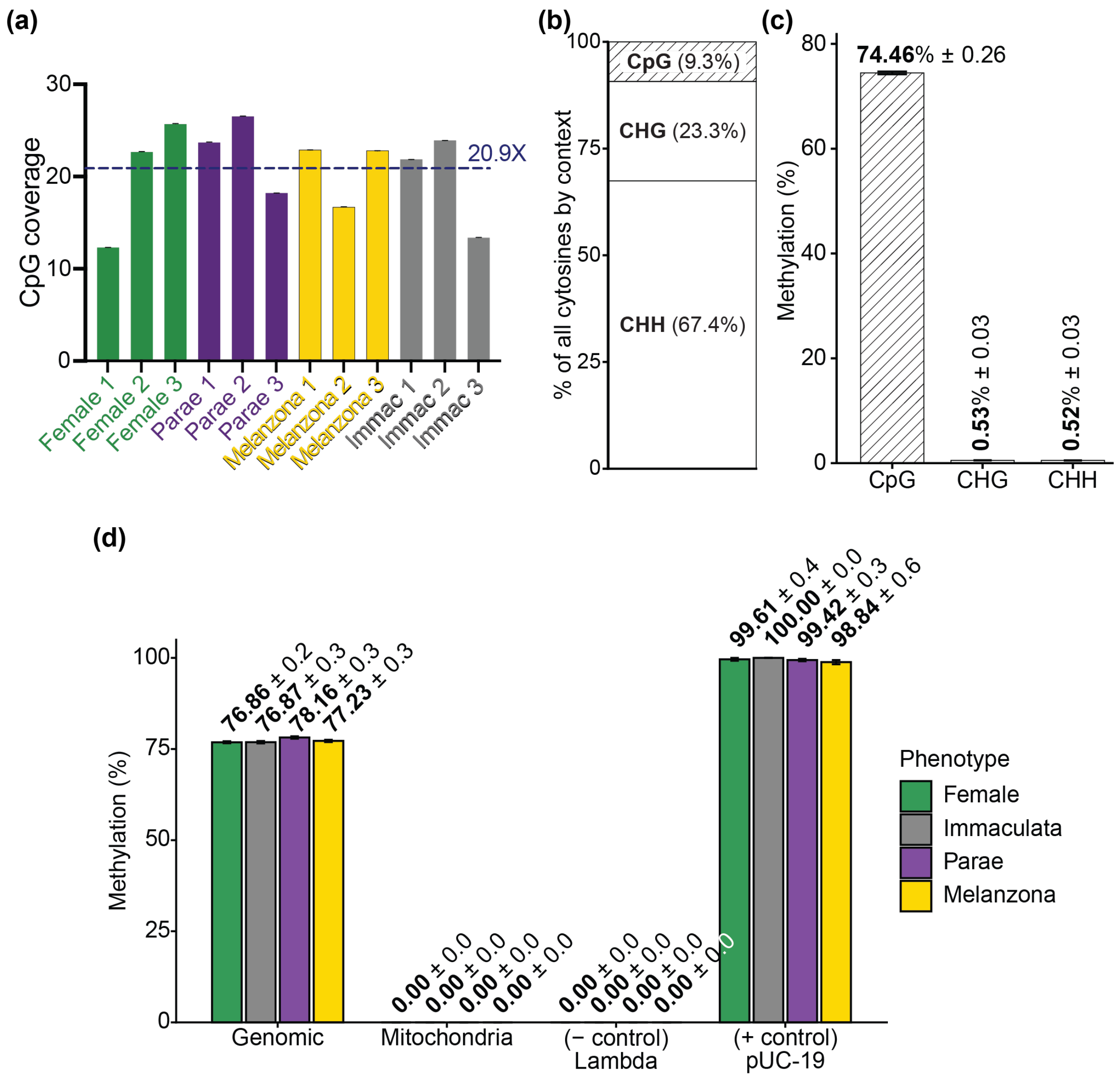


**Supplementary Figure 1.** Quality control of EM-seq libraries shows near-complete conversion and methylation restricted to CpG sites. (a) CpG coverage of each sample (mean ± SEM); the dashed line shows the mean across samples (20.9X). (b) Proportion of all sequenced cytosines that occur in CpG, CHG, and CHH contexts. (c) Methylation level of all sequenced cytosines in each context (mean ± 1 SEM, n = 12), showing that methylation is largely restricted to CpG sites. (d) CpG methylation of genomic DNA, mitochondrial DNA, and two spike-in controls in each phenotype (mean ± 1 SEM across the 3 samples of each phenotype): unmethylated lambda DNA, whose near-zero methylation indicates near-complete conversion, and CpG-methylated pUC19, whose near-complete methylation indicates that methylated cytosines were protected from conversion. Genomic values differ slightly between (c) and (d) because they were calculated with Bismark and DNMTools, respectively.

Alt text: Four panels of bar charts summarizing EM-seq data quality. Panel a shows mean CpG coverage for each of the 12 samples, ranging from about 12× to 27×, with a dashed line at the overall mean of 20.9×. Panel b is a stacked bar showing that 9.3% of sequenced cytosines are in CpG context, 23.3% in CHG, and 67.4% in CHH. Panel c shows methylation of 74.5% at CpG sites and about 0.5% at CHG and CHH sites. Panel d shows CpG methylation for each phenotype: about 77–78% for genomic DNA, 0% for mitochondrial DNA and the unmethylated lambda control, and 98.8–100% for the methylated pUC19 control.

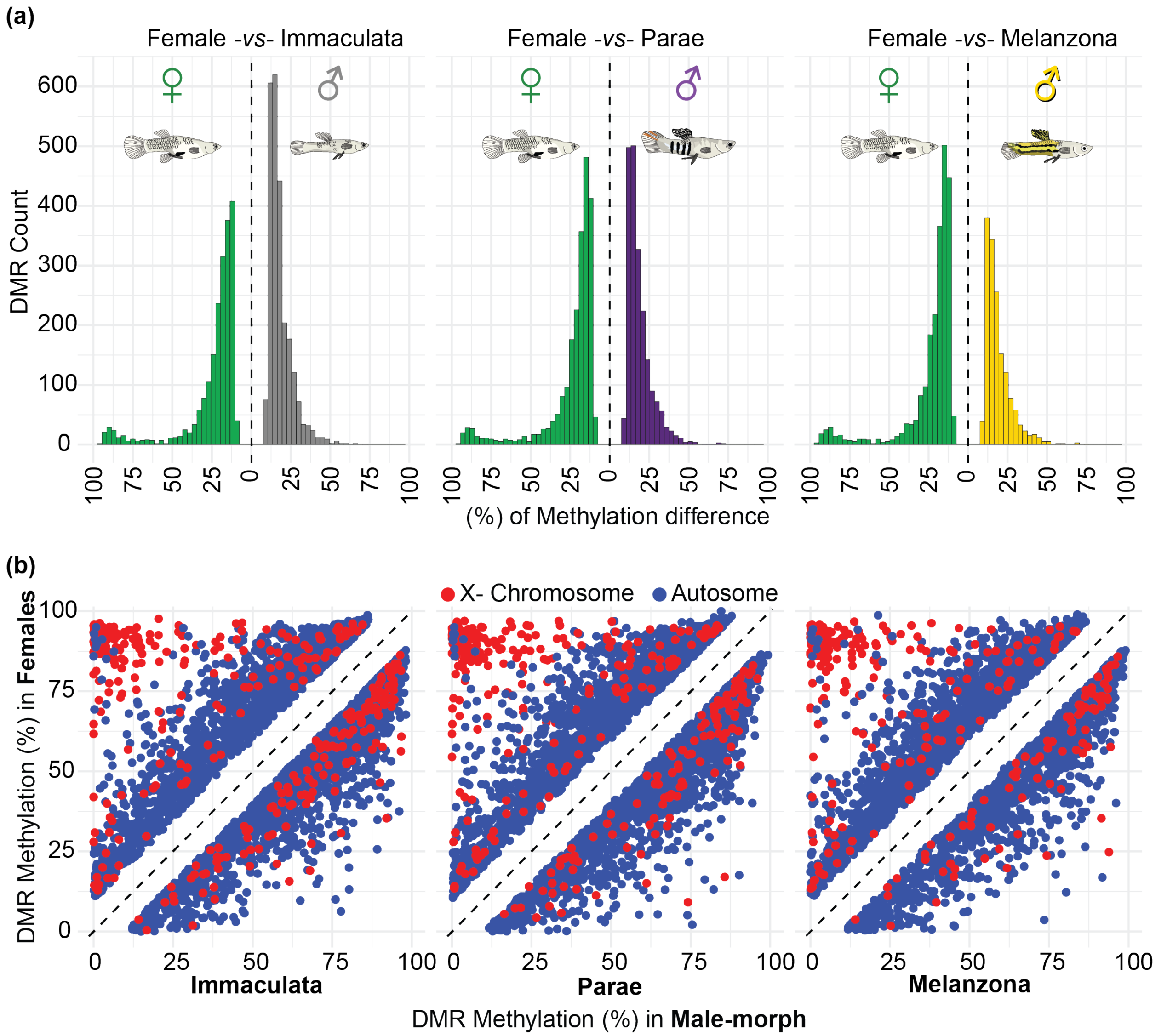


**Supplementary Figure 2.** Most female-male DMRs show modest differences in either direction, but a subset of X-linked DMRs are highly methylated in females and nearly unmethylated in males. (a) Distribution of methylation differences for all DMRs between females and each male morph (female vs. Immaculata, n = 4,496; female vs. Parae, n = 4,289; female vs. Melanzona, n = 3,812). DMRs more highly methylated in females are shown to the left of zero and those more highly methylated in males to the right; differences below 10% are absent because of the DMR detection threshold. (b) Mean methylation of each DMR in females versus each male morph, for autosomal (blue) and X-linked (red) DMRs; points above the dashed diagonal are more highly methylated in females.

Alt text: Panel a shows three histograms, one for females compared with each male morph, of the number of differentially methylated regions by size of methylation difference, with regions more methylated in females to the left of zero and those more methylated in males to the right. Most differences fall between 10% and 30% in both directions, but each female-side histogram has a small second peak near 85–100%. Panel b shows three scatterplots of each region's methylation in females (y-axis) against a male morph (x-axis), with autosomal regions in blue and X-linked regions in red. Points fall in two bands parallel to the diagonal, and a cluster of red X-linked points sits in the upper left, near 90% methylation in females and near 0% in males.

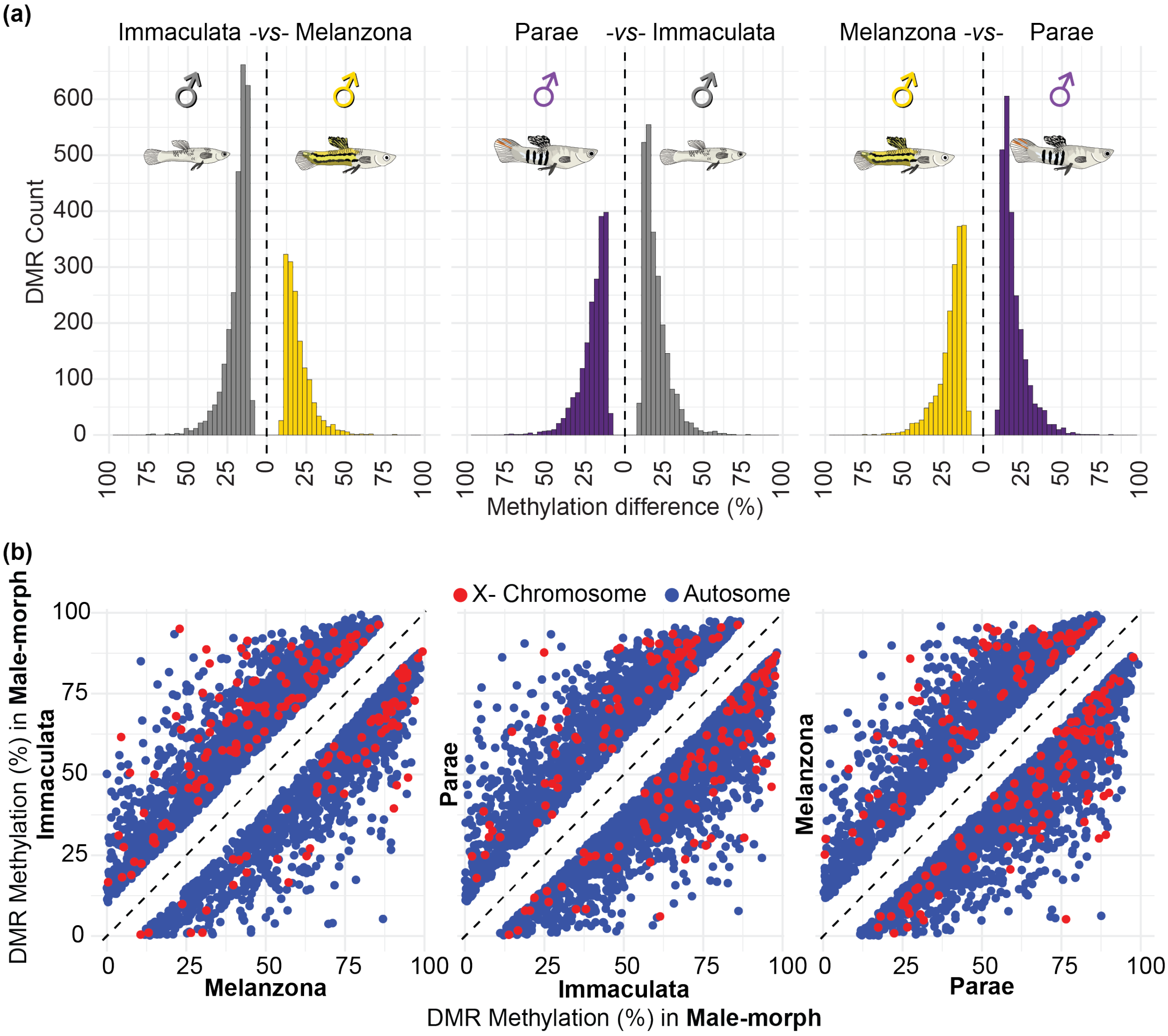


**Supplementary Figure 3.** DMRs between male morphs show modest methylation differences in either direction, with no X-linked cluster comparable to that between the sexes. (a) Distribution of methylation differences for all DMRs between each pair of male morphs (Immaculata vs. Melanzona, n = 4,142; Parae vs. Immaculata, n = 4,334; Melanzona vs. Parae, n = 4,278). DMRs more highly methylated in the first-named morph are shown to the left of zero and those more highly methylated in the second to the right; differences below 10% are absent because of the DMR detection threshold. (b) Mean methylation of each DMR in the first-named (y-axis) versus second-named (x-axis) morph, for autosomal (blue) and X-linked (red) DMRs; points above the dashed diagonal are more highly methylated in the first-named morph.

Alt text: Panel a shows three histograms, one for each pair of male morphs, of the number of differentially methylated regions by size of methylation difference, with regions more methylated in the first-named morph to the left of zero. Nearly all differences fall between 10% and 40%, with no second peak at large differences. In each pair, one morph has more regions with higher methylation: Immaculata relative to both other morphs, and Parae relative to Melanzona. Panel b shows three scatterplots of each region's methylation in one morph against another, with autosomal regions in blue and X-linked regions in red. Points fall in two bands parallel to the diagonal, and X-linked points are mixed among autosomal points with no separate cluster.

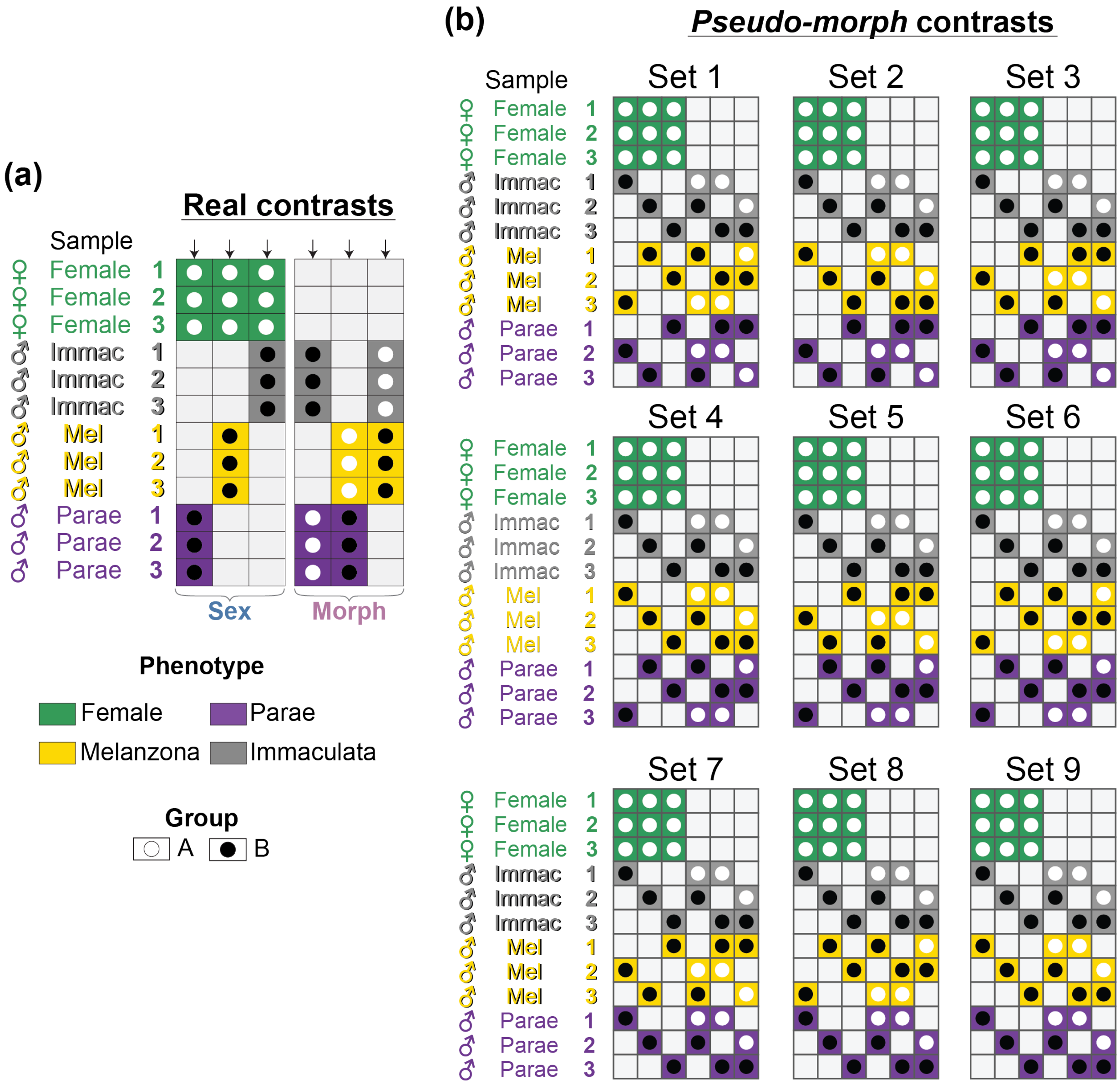


**Supplementary Figure 4**. Design and group composition of the real and randomized (pseudo-morph) comparisons. Each column is one pairwise comparison and each row one individual; open and filled circles mark the two groups compared, cell color indicates phenotype, and empty cells indicate individuals not included in that comparison. (a) The six comparisons among the real groups (females and the three male morphs). (b) The nine randomized sets, each containing the same six comparisons as in (a) but with the nine males reassigned to three pseudo-morphs, each containing one male of each morph. Labels identify the three pseudo-morphs in each set (e.g., 01_02_03 = pseudo-morphs 1–3), matching the identifiers in Supplementary Table 3.

Alt text: Grid diagrams showing which individuals were compared in each analysis. Rows are the 12 individuals (three females and three each of Immaculata, Melanzona, and Parae males), and columns are pairwise comparisons; colored cells with open or filled circles mark the two groups in each comparison. Panel a shows the six real comparisons: females against each male morph, and each pair of male morphs. Panel b shows nine smaller grids, one per randomized set, each with the same six-comparison layout, but with each male group (pseudo-morph) made up of one Immaculata, one Melanzona, and one Parae male.

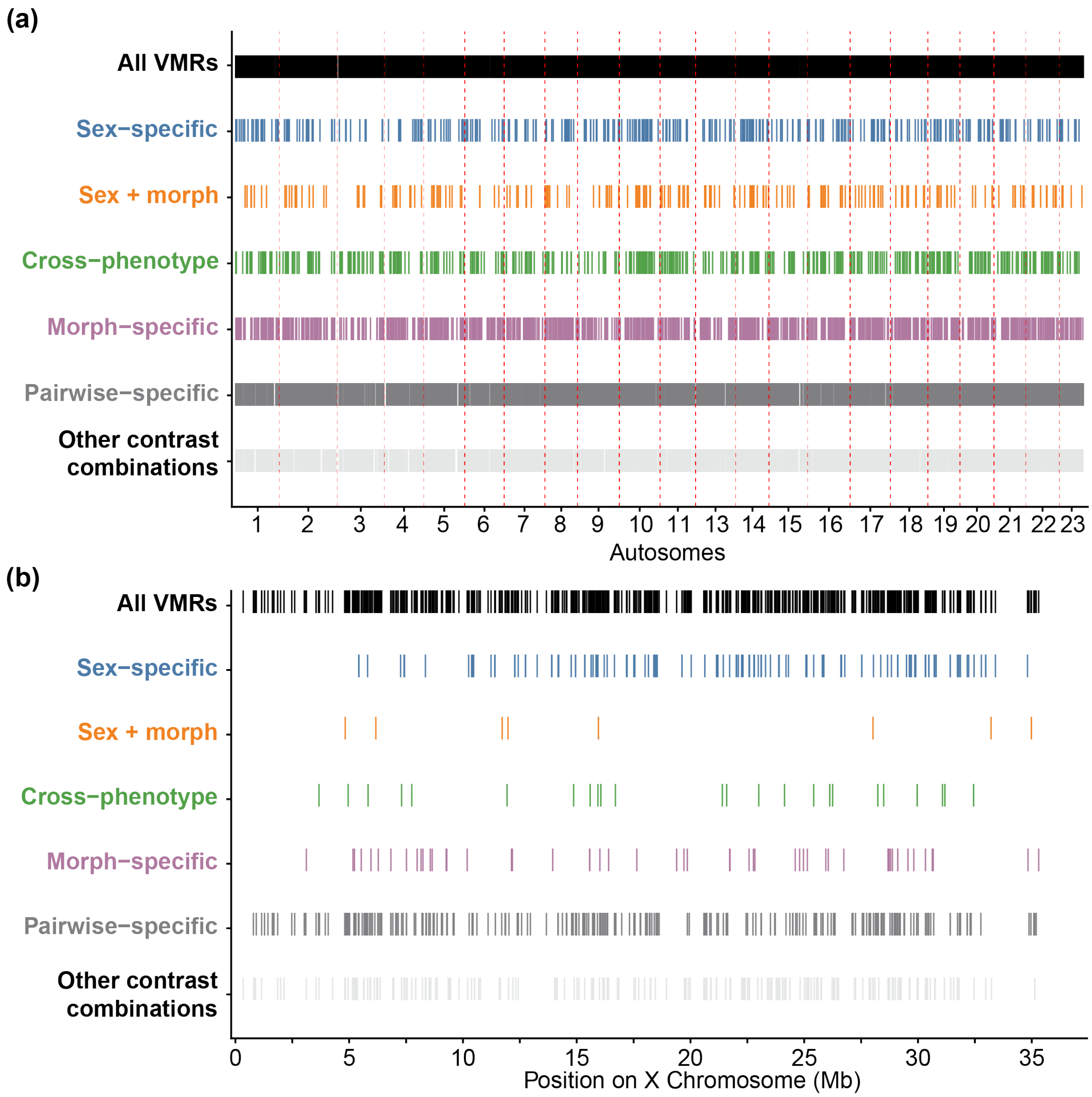


**Supplementary Figure 5.** VMRs of every category are distributed throughout the genome. Positions of VMRs on (a) the autosomes, shown end to end in chromosome order (chromosome 12, the X, is shown separately), and (b) the X chromosome, for all VMRs and for each category defined in Figure 2. Each tick marks one VMR; red dashed lines in (a) mark chromosome boundaries.

Alt text: Two genome maps in which each vertical tick marks one variably methylated region, with separate tracks for all regions and for each category (sex-specific, sex and morph, cross-phenotype, morph-specific, pairwise-specific, and other). Panel a places the autosomes end to end, separated by dashed lines; ticks for every category are spread along all autosomes, with no clusters in particular chromosomes. Panel b shows the X chromosome from 0 to about 37 Mb; sex-specific, morph-specific, and pairwise-specific regions occur along most of its length, while sex and morph and cross-phenotype regions are sparse.

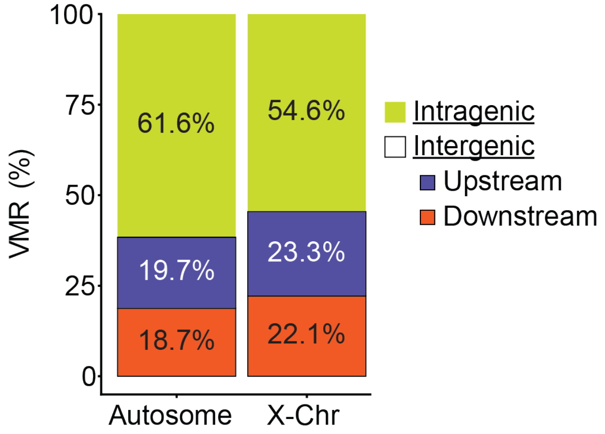


| partition | n chr | observed intragenic | observed intergenic | observed total | expected intragenic | expected intergenic | $X^2$ | df | pvalue |
|---|---|---|---|---|---|---|---|---|---|
| **AUTO** | 22 | 6731 | 4192 | 10923 | 6379.0 | 4544.0 | 100.95 | 22 | 4.399E-12 |
| **WG** | 23 | 7118 | 4514 | 11632 | 6748.1 | 4883.9 | 102.76 | 23 | 4.684E-12 |

**Supplementary Figure 6.** VMRs overlap genes more often than expected by chance. Bars show the percentage of VMRs on the autosomes and X chromosome that overlap a gene (intragenic) or lie upstream/downstream of the nearest gene, defined by gene orientation (intergenic). The table gives observed and expected numbers of intragenic and intergenic VMRs for the autosomes (AUTO) and whole genome (WG), with expected values based on the proportion of each chromosome occupied by genes. $\chi^2$ tests compare observed and expected counts across chromosomes; more VMRs were intragenic than expected in both partitions.

Alt text: Stacked bar chart and table describing where variably methylated regions fall relative to genes. On the autosomes, 61.6% of regions overlap a gene, 19.7% are upstream, and 18.7% are downstream of the nearest gene; on the X chromosome, the values are 54.6%, 23.3%, and 22.1%. The table below lists observed and expected numbers of regions inside and outside genes: 6,731 observed versus 6,379 expected inside genes on the autosomes, and 7,118 versus 6,748 genome-wide, with chi-squared tests significant in both cases ($p < 10^{-11}$).

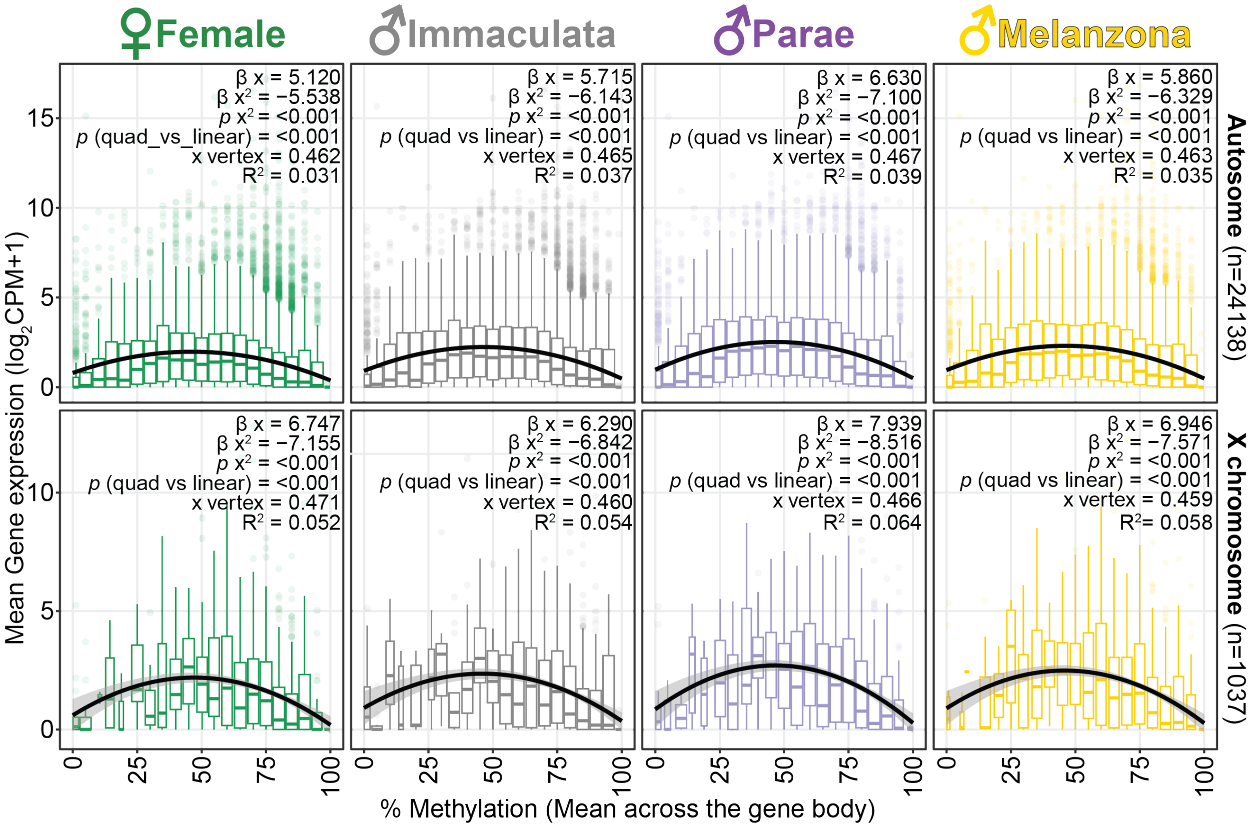


**Supplementary Figure 7.** Gene expression is highest at intermediate gene-body methylation in all phenotypes. For each phenotype, mean expression of each gene ($\log_2$CPM + 1) is plotted against its mean gene-body methylation, on the autosomes (top) and X chromosome (bottom); expression was measured in separate individuals from the same population (see Methods). Boxplots summarize genes in bins of methylation. Black lines show quadratic fits with 95% confidence intervals; each panel gives the linear (β x) and quadratic (β $x^2$) coefficients, the methylation level at peak expression (x vertex), $R^2$, and tests of the quadratic term and of the quadratic versus linear model. Models were fit with methylation as a proportion (0-1), so coefficients and vertices are on that scale.

Alt text: Eight scatter-and-boxplot panels, one per phenotype (female, Immaculata, Parae, Melanzona) for the autosomes (top row) and the X chromosome (bottom row), plotting mean gene expression against mean gene-body methylation from 0% to 100%. Boxplots summarize genes in methylation bins, and a black curved line shows a quadratic fit. In every panel, expression is lowest at very low and very high methylation and peaks near 46–47% methylation, forming an arch. The fitted curves explain 3–6% of the variation in expression.

(a)

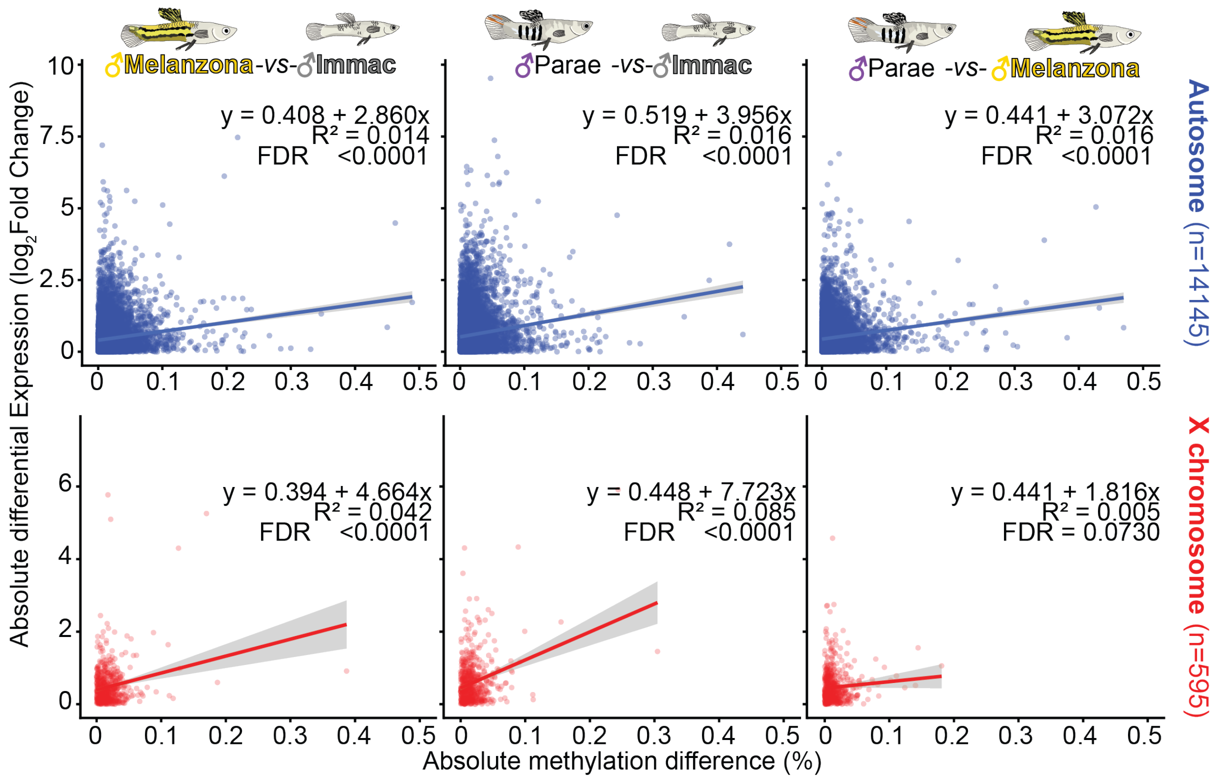

♂Melanzona-vs-♂Immac
♂Parae -vs-♂Immac
♂Parae -vs- ♂Melanzona
Absolute differential Expression ($\log_2$Fold Change)
y = 0.408 + 2.860x
R² = 0.014
FDR <0.0001
y = 0.519 + 3.956x
R² = 0.016
FDR <0.0001
y = 0.441 + 3.072x
R² = 0.016
FDR <0.0001
Autosome (n=14145)
y = 0.394 + 4.664x
R² = 0.042
FDR <0.0001
y = 0.448 + 7.723x
R² = 0.085
FDR <0.0001
y = 0.441 + 1.816x
R² = 0.005
FDR = 0.0730
X chromosome (n=595)
Absolute methylation difference (%)


(b)

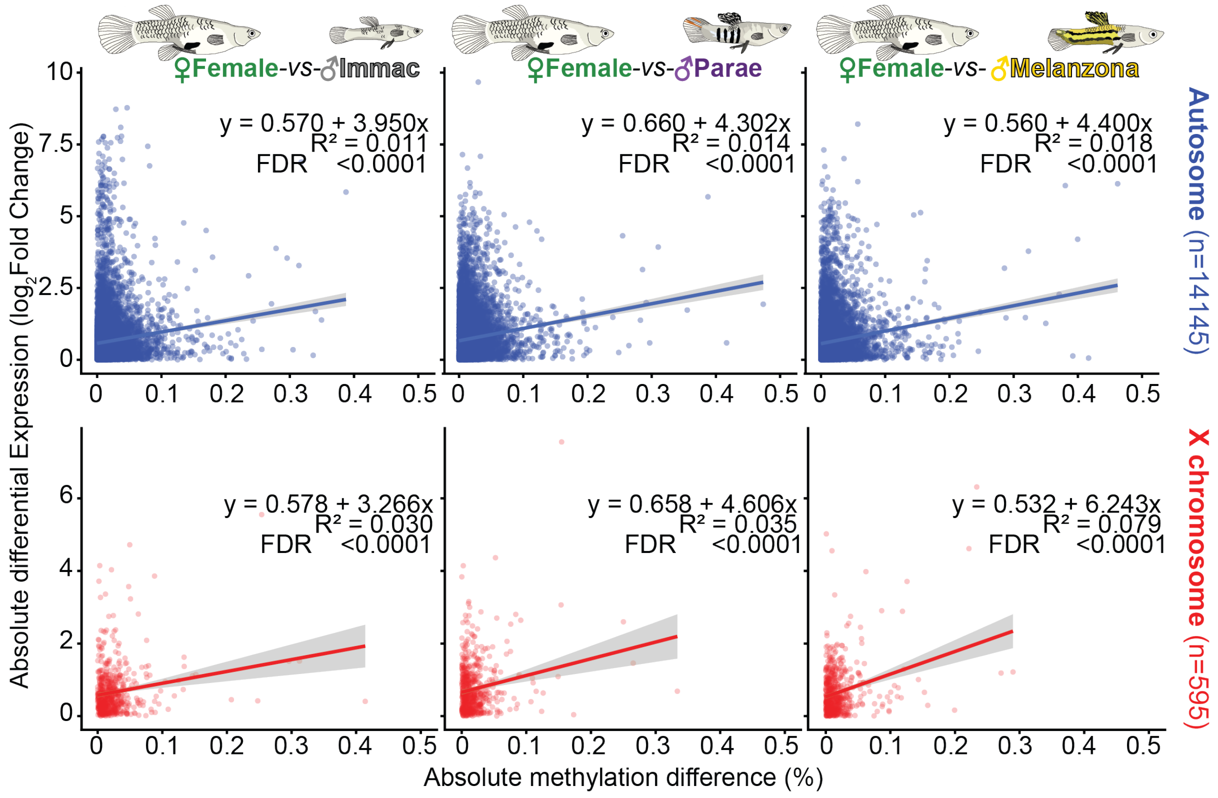

♀Female-vs-♂Immac
♀Female-vs-♂Parae
♀Female-vs- ♂Melanzona
Absolute differential Expression ($\log_2$Fold Change)
y = 0.570 + 3.950x
R² = 0.011
FDR <0.0001
y = 0.660 + 4.302x
R² = 0.014
FDR <0.0001
y = 0.560 + 4.400x
R² = 0.018
FDR <0.0001
Autosome (n=14145)
y = 0.578 + 3.266x
R² = 0.030
FDR <0.0001
y = 0.658 + 4.606x
R² = 0.035
FDR <0.0001
y = 0.532 + 6.243x
R² = 0.079
FDR <0.0001
X chromosome (n=595)
Absolute methylation difference (%)

**Supplementary Figure 8.** Methylation differences between phenotypes are weakly but consistently associated with expression differences. For each gene, the absolute $\log_2$ fold change in expression is plotted against the absolute difference in gene-body methylation between (a) pairs of male morphs and (b) females and each male morph, on the autosomes (blue) and X chromosome (red); expression was measured in separate individuals from the same population (see Methods). Lines show linear fits with 95% confidence intervals; each panel gives the fitted equation, $R^2$, and Benjamini–Hochberg-adjusted p-value. Models were fit with methylation as a proportion (0-1), so slopes are on that scale.

Alt text: Twelve scatterplots of the absolute difference in gene expression ($\log_2$ fold change) against the absolute difference in gene-body methylation, one for each of six phenotype comparisons on the autosomes (blue) and X chromosome (red). Panel a shows the three male-morph comparisons and panel b shows females compared with each male morph. Most genes cluster near zero methylation difference with a wide spread of expression differences. Each panel has a straight fitted line with a shallow positive slope; the lines explain 1–2% of the variation on the autosomes and up to about 8% on the X chromosome. All slopes are positive, and all are significant except the X-chromosome comparison of Parae and Melanzona.
